# Inverted colored de Bruijn Graph for practical kmer sets storage

**DOI:** 10.64898/2025.12.08.692073

**Authors:** Timothé Rouzé, Rayan Chikhi, Antoine Limasset

## Abstract

Petabases of sequencing data in the Sequence Read Archive (SRA) present a significant challenge for holistic reanalysis due to their sheer volume. Recent efforts have assembled this data into terabytes of unitigs, an efficient *k*-mer set representation that can reduce data size by an order of magnitude. However, these unitigs were compressed on a per-accession basis, leaving substantial cross-sample redundancy unexploited. While co-compression of related samples offers high space-saving potential, existing tools lack targeted decompression: the ability to retrieve specific documents at a cost proportional to their individual sizes rather than that of the entire collection.

This paper introduces the “inverted de Bruijn graph” property, formalizing the concept of efficient targeted decompression, and presents kloe, its first implementation. kloe is a compression method for large, highly similar *k*-mer multi-sets, such as collections of unitigs from related samples. Unlike existing approaches that map *k*-mers to colors (samples), kloe takes a complementary route by performing color-to-*k*-mer mapping, associating samples with their respective *k*-mer sets. This enables targeted decompression of any chosen sample’s *k*-mer content. At its core, kloe utilizes a new sequence construct called “monochromatigs,” drawing on concepts from simplitigs and monotigs to achieve both significant space savings and efficient retrieval. Finally, a central aim of this work is to highlight this novel problem area, which we argue is critically understudied compared to colored de Bruijn graphs. The associated tool is available as an open source project at github.com/TimRouze/KLOE

## 1 Introduction

kmers have become a cornerstone of efficient biological sequence analysis. Unitigs [5], which represent sets of overlapping *k*-mers as contiguous sequences, are a widely used and efficient representation for *k*-mer sets across many applications [4, 6, 9]. This approach has been so successful that the Logan [8] project now provides access to the all accession of the Sequence Read Archive (SRA) transformed into unitigs. This initiative processed 50 petabases of raw SRA data, resulting in approximately 2.18 petabytes of unitig sequences. However, this very success highlights a significant compression challenge. Crucially, Logan unitigs are compressed on a per-accession basis using the `zstd` compressor. While this strategy substantially reduces storage requirements, it fails to exploit the significant inter-sample redundancy found in the Sequence Read Archive (SRA) which contain, for example, hundreds of thousands of human sequencing experiments, leading to considerable content overlap. This is illustrated in Figure 1, where more than 2 billion *k*-mers in a subset of human Logan WGS samples are observed in all 128 analyzed files. Therefore, we hypothesize that by leveraging this pervasive redundancy across the entire SRA, as represented in the Logan database, orders of magnitude improvements in compression can be achieved.

**Figure 1.**
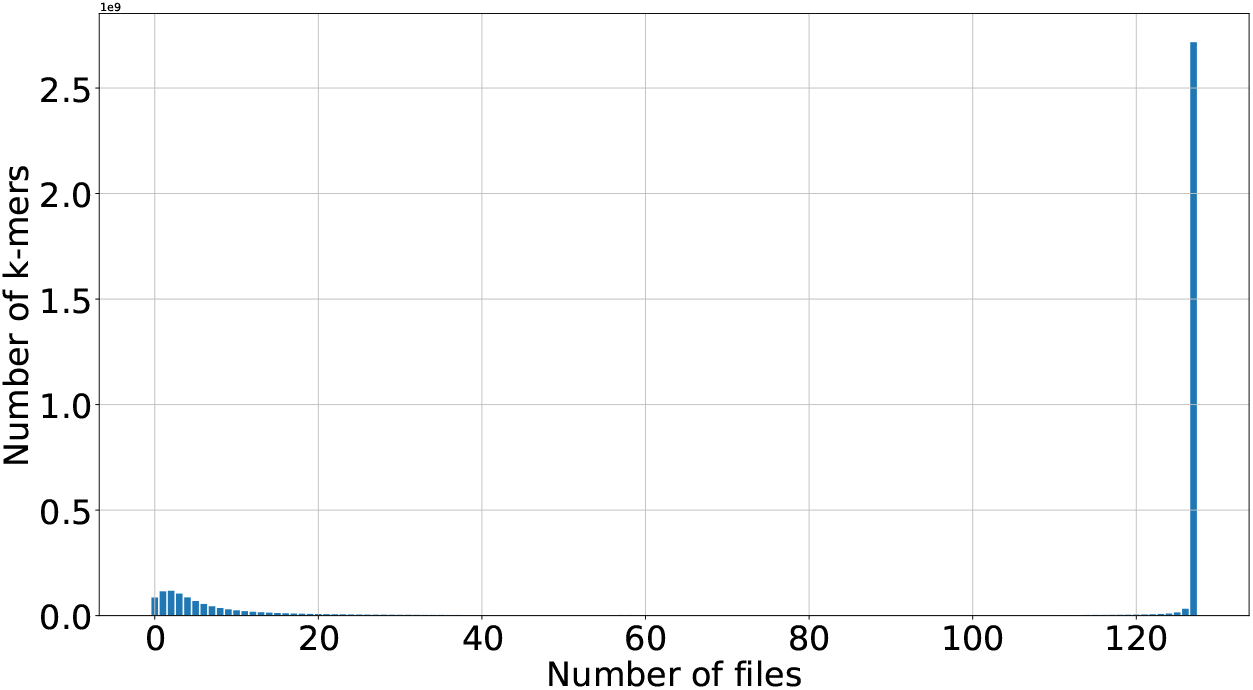
Bar plot showing the number of *k*-mers (y axis) that are seen in human unitig files (x axis, 128 files total) for a subset of human Logan WGS samples. Each bar of height *h* at coordinate *x* corresponds to the number *h* of *k*-mers that are found in exactly *x* of the files. Here, the bar at position *y* = 128 shows that a majority of *k*-mers (over 2.5 billion) are seen in all files, representing a formidable opportunity for compression.

However, compressing large biological sequence collections into monolithic archives, while maximizing potential compression by exploiting inter-dataset redundancies, creates a significant hurdle: entire archives must be decompressed to retrieve even single samples. This inefficiency becomes critical as public DNA repositories grow exponentially, rendering full decompression impractical for routine access. Consequently, there is a pressing need to develop methods for efficient, direct decompression of targeted subsets from these large compressed collections, instead of fragmenting them into less efficient smaller units to improve usability. For instance, in the context of genome collections, `agc` [10] pioneered this kind of functionality, allowing for the targeted retrieval of individual genomes from a compressed collection. However, `agc`‘s alignment-based mechanisms, optimized for relatively complete genome sequences, are not directly transferable to unitig collections. In practice, `agc` is not able to compress them. Being generally shorter, far more numerous, and more fragmented than assembled genomes, they pose significant challenges for such alignment-based strategies. Other genome collection compressors, like Miniphy [3], also typically lack this targeted retrieval capability, necessitating complete or large-batch decompression. Furthermore, tools designed to compress raw sequencing datasets [12], specialized for particular read properties (such as length or redundancy distribution), are unsuitable for this context. Consequently, a solution specifically designed for the properties of unitig collections, offering efficient targeted decompression, is currently lacking.

Colored de Bruijn graphs (cDBGs), the subject of extensive recent research [17], are natural candidates for indexing collections of *k*-mers sets. Their primary function is efficiently retrieving the origin sets for a query *k*-mer *k*_*q*_ from a collection *S* = {*s*_1_, … , *s*_*n*_ }, i.e., determining *q*(*k*_*q*_) = {*s*_*i*_ ∈ *S* : *k*_*q*_ ∈ *s*_*i*_ }. Numerous lightweight techniques, either exact or probabilistic (the latter potentially yielding false positives for some *k*-mers), have been proposed for this task. For example, `metagraph` [14] uses annotated de Bruijn graphs (colors) for efficient *k*-mer presence queries, while `ggcat` [9] rapidly constructs compressed cDBGs from large *k*-mer sets. Other prominent cDBG indexes include `fulgor` [11], which leverages Minimal Perfect Hash Functions [19], and `themisto` [1], based on the Spectral Burrows Wheeler Transform. As static structures encoding both sequence data and their sources, cDBGs also serve as *k*-mer set compression tools; ESS-color [21], focuses on compressing colored de Bruijn graphs by mapping a spectrum-preserving string set (SPSS) [2, 20] to a color matrix using SSHash [19]. A compressor by Kitaya & Shibuya also leverages DBGs to perform compression leveraging local textual similarities from multiple *k*-mer sets [16].

cDBG indexing tools that perform lossless *k*-mer indexing theoretically permit reconstruction of the original *k*-mer sets. Though not their primary design goal, these indexing tools often achieve compact cDBG representations, making them space-efficient. A key limitation, however, is that they are optimized towards *k*-mer -to-color queries. Our objective is the inverse: efficiently retrieving all *k*-mers originating from a specific input dataset *s*_*j*_ (or a set of input datasets). With cDBGs, recovering all *k*-mers from *s*_*j*_ is possible in theory. However, to the best of our knowledge this feature has never been implemented. In [17] data structure for collections of *k*-mers are separated in two groups. The first group dubbed “color-aggregative” comprises structures that offer *k*-mer to color organization. For those, a natural solution would be to iterate over every *k*-mer in the graph and check their color-sets, which would lead to a complexity in *O*(|∪*S* |). The second group, called “*k*-mer -aggregative” represents structures that allow iteration over each color-set, implying a complexity in *O*(|*Z* |) with *Z* = *z*1, … , *z*_*x*_ the set of existing color sets. These two solutions imply a substantial overhead compared to an ideal complexity in the order of *O*(|*s*_*j*_ |).

Therefore, the primary goal of this paper is to define data structures capable of compressing large collections of unitigs while simultaneously enabling efficient random access to any given subset of unitigs, a functionality currently lacking in existing approaches. A second objective is to introduce `kloe`, a first proof-of-concept embodying these principles. Finally, we aim to stimulate the research community’s interest in this pressing issue, which addresses tangible and challenging applicative needs.

## 2 Methods

This section begins by establishing the necessary definitions used throughout our work. Subsequently, we detail the input data characteristics utilized in practice by our `kloe` compressor. Following this, we describe the compression mechanism itself, and finally, we outline the corresponding decompression mechanism.

## 2.1 Key definitions

Here, we introduce key terminology that underlies our work, starting with *k*-mers and de Bruijn graph.

### Definition 1 (*k*-mers)

*A k****-mer*** *is a string of fixed length k >* 0 *over the alphabet* Σ = {*A, T, C, G* }. *The set of k-mers of a sequence S, denoted K*(*S*), *is the set of all substrings of length k that can be extracted from S. Similarly, the set of k-mers of a set of sequences* 𝒮, *denoted K* (𝒮), *is the union of all k-mers extracted from each sequence in* 𝒮 *(i*.*e*., *K*(𝒮) = ∪ _*s*∈ 𝒮_*K*(*s*)*). For simplicity in these definitions, reverse complements are often omitted from explicit notation. However, in many implementations, each k-mer is represented by its canonical form, which is the lexicographically smaller of the k-mer and its reverse complement. Subsequent definitions assume k-mers can be handled in their canonical form where appropriate*.

### Definition 2 (de Bruijn Graph (dbg))

*Given a set of input sequences* _*in*_ *and an integer k >* 0, *a* ***de Bruijn graph (dbg)***, *denoted G*_*k*_(𝒮_*in*_) = (*V, E*), *is a directed graph. The set of vertices V consists of all distinct k-mers (potentially in their canonical form) extracted from the sequences in* 𝒮_*in*_; *thus, V* = *K*(𝒮_*in*_). *The set of edges E contains a directed edge* (*u, v*) *from k-mer u* ∈ *V to k-mer v* ∈ *V if and only if the* (*k* − 1)*-length suffix of u is identical to the* (*k* − 1)*-length prefix of v. This definition is* ***node-centric***, *as the edges E are implicitly defined by the vertex set V* . *The graph is therefore informationally equivalent to its set of k-mers V [4]*.

When handling multiple *k*-mer sets, a common approach is to consolidate their de Bruijn graphs into a single structure dubbed colored de Bruijn graph [13]. This allows for querying a *k*-mer to determine its “colors”, defining its presence in the different datasets.

### Definition 3 (Colored de Bruijn Graph (cdbg))

*Let* ℒ _𝒮_ = (𝒮_1_, 𝒮_2_, … , 𝒮_*m*_) *be a list of m sets of sequences. A* ***cdbg*** *corresponding to* ℒ_𝒮_ *is defined based on a consistent k-mer length k is a list of m de Bruijn graphs*, 𝒢_*collection*_ = (*G*_1_, *G*_2_, … , *G*_*m*_). *Each graph G*_*i*_ = *G*_*k*_(𝒮_*i*_) *is the de Bruijn graph constructed from the set of sequences* 𝒮_*i*_. *The set of all k-mers present in the cdbg is* 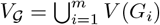, *where V* (*G*_*i*_) *is the vertex set of G*_*i*_. *This collection of graphs is called colored because, for each* k*-mer* , *it knows the de Bruijn graph(s) of origin*.

### Definition 4 (Color-set of a *k*-mer)

*Given a cdbg built from m sets of sequences, leading to m component de Bruijn graphs* (*G*_1_, *G*_2_, … , *G*_*m*_), *the* ***color-set*** *of a k-mer x (of the same length k) is a boolean vector C*(*x*) = (*c*_1_, *c*_2_, … , *c*_*m*_). *Each element c*_*i*_ *is true if the k-mer x is present as a node in the i-th component de Bruijn graph G*_*i*_ *(i*.*e*., *x* ∈ *V* (*G*_*i*_)*), and false otherwise*.

To manage the often extensive size of (colored) de Bruijn graphs, compaction techniques exploiting the fact that *k*-mers overlap can be applied to represent *k*-mers sets as sequences in a space efficient manner [5]. These techniques known as spectrum preserving string sets [22], [2] cover unitigs, simplitigs [2], matchtigs [24] or eurlertig [23] and were generalized as masked superstrings [25].

### Definition 5 (Spectrum Preserving String Set for a cdbg)

*Let a cdbg be defined from m component graphs* (*G*_1_, … , *G*_*m*_), *where V* (*G*_*i*_) *is the set of k-mer nodes for each G*_*i*_. *A* ***Spectrum Preserving String Set (SPSS)*** *for this cdbg is a set of strings*, 𝒮_*cdbg*_, *such that its derived set of k-mers, K*(𝒮_*cdbg*_) *(using K*(·)*), is equal to the union of the k-mer sets of all component graphs. Thus*, 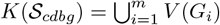.

Building from the previous concepts, we introduce a specific spectrum preserving string set scheme for colored de Bruijn graphs which we call monochromatigs. Monochromatigs are conceptually similar to *monotigs* [18]; however, the defining constraint for Monochromatigs is based on the uniformity of *k*-mer presence/absence patterns across datasets (their color set), rather than *k*-mer abundances.

### Definition 6 (Monochromatigs)

*A set of sequences* ℳ *is defined as* ***Monochromatigs*** *with respect to a cdbg if it is a Spectrum Preserving String Set (SPSS) for that cdbg, and additionally, every sequence s* ∈ ℳ *is* ***monochromatic***. *A sequence s is considered monochromatic if all k-mers that can be extracted from s (i*.*e*., *all x* ∈ *K*({*s* })*) possess the exact same color set C*(*x*). *This means that for any given sequence in* ℳ, *all its constituent k-mers are present in the same combination of original datasets*.

Such representation has been used in several indexes such as `themisto`, `fulgor` and `ggcat` [1], [19], [9].

We will define the query capabilities and performance standards for an efficient colored de Bruijn graph, along with the desirable properties of an inverted index designed for the inverse query, which retrieves all *k*-mers associated with a given set of colors (see Figure 2).

**Figure 2.**
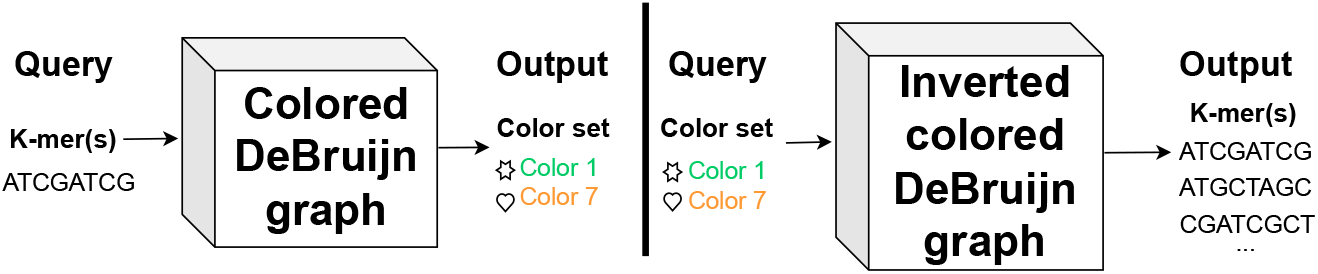
Comparison between *k*-mer membership queries (colored DBG index) with targeted *k*-mer extraction queries (inverted colored DBG index)

### Definition 7 (cdbg Index)

*A* ***cdbg Index*** *is a data structure associated with a cdbg (effectively built from m component datasets). For any given k-mer x, it is able to return its color set C*(*x*) = (*c*_1_, *c*_2_, … , *c*_*m*_), *which indicates the presence (true) or absence (false) of x in each of the m components, most implementation offering a query time of O*(|*m*|).

We can now formulate the main problem.

### Problem 1 (Targeted decompression)

*Given a collection* ℒ_𝒮_ = (𝒮_1_, 𝒮_2_, … , 𝒮_*m*_) *of sets of sequences, compress* ℒ_𝒮_ *into a binary representation B so that each S*_*j*_ *can be extracted from B in O*(|*S*_*j*_|).

To the best of our knowledge, there is currently no practical solution to Problem 1. To reach *O*(|𝒮_*j*_|), here we propose a novel data structure called Inverted colored de Bruijn graph.

### Definition 8 (Inverted cdbg Index)

*An* ***Inverted cdbg Index*** *is a data structure associated with a cdbg (effectively built from m component datasets). For a given input color set* 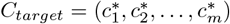 *it is able to return the set* 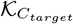 *of all k-mers* {*x* } *from the cdbg whose color set C*(*x*) *is equal to C*_*target*_, *with a query time of* 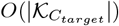, *where* 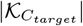 *is the size of the returned set*.

This allows us to formally state our result. Problem 1 can be solved by building a cdbg of ℒ_𝒮_ then an inverted cdbg index for the cdbg.

## 2.2 Input data

The `kloe` compressor is specifically designed to compress various types of spectrum-preserving string sets (SPSS), such as unitigs [5], simplitigs [2], eulertigs [23], and matchtigs [24]. It can process files produced by assemblers like `bcalm2` [7], `ggcat` [9], and `Cuttlefish` [15], as well as those from collections like such as Logan unitigs or contigs [8]. Any type of dataset containing samples from different sources that features strong inter-sample redundancy is a good candidate for compression using the method presented in this paper.

## 2.3 Compression method

We present `kloe`, a proof-of-concept tool that implements an inverted de Bruijn graph index. `kloe` computes monochromatigs from input files and organizes them with metadata to create this inverted index, enabling targeted decompression of DNA sequences in *O*(|*s*_*j*_ |). The compression workflow employed by `kloe` is illustrated in Figure 3.

**Figure 3.**
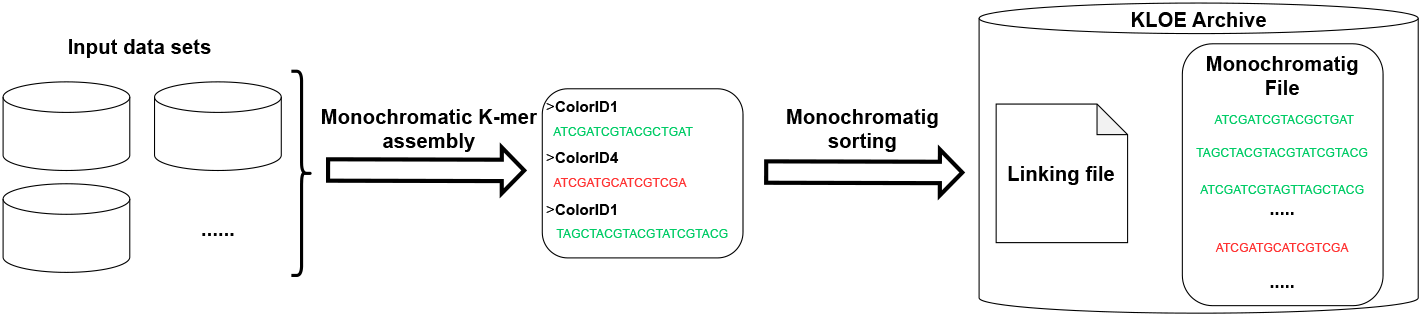
Illustration of the compression workflow in `kloe`. **Monochromatic *k*-mer assembly**: *k*-mers are assembled as monochromatigs and stored temporarily before the sorting phase. **Monochromatig sorting**: Monochromatigs are retrieved from the temporary file and written to the final output in sorted order.

### 2.3.1 Monochromatig generation

Initially, `kloe` assembles *k*-mers into monochromatigs. This assembly mimics the greedy algorithm presented in [2]. The original greedy algorithm traverses the *k*-mer graph, assembling the first encountered adjacent *k*-mers not yet part of a simplitig, continuing until no unused neighboring *k*-mers remain. We augment this by requiring assembled *k*-mers to share the same color set to form a monochromatig. While this typically results in shorter simplitigs, it enables targeted decompression by allowing sequences to be grouped by their color sets, as detailed later.

### 2.3.2 Sorting monochromatigs

During monochromatig generation, their sequences are streamed to a temporary file. Once all *k*-mers are assembled, these monochromatigs are read from the temporary file and sorted by their color set. The final monochromatig file therefore consists of a succession of color *buckets*, where each bucket contains monochromatigs sharing the same color set. This organization minimizes the disk access required to retrieve all *k*-mers of interest during targeted decompression. As depicted in Figure 3, this sorting into color buckets streamlines targeted decompression and can reduce disk operations by allowing direct access to relevant monochromatig groups rather than processing them individually.

### 2.3.3 Linking

Upon completion of the sorting operation, a linking file containing necessary metadata is created. This file maps each color bucket to:

- a bitvector representing its color set,
- the starting position of the bucket within the monochromatig file,
- a list of integers detailing the sizes of each monochromatig in that bucket.

Currently, this linking file is compressed using `zstd` (with the -12 option) but is not yet fully optimized for minimal compressibility. Further optimizations could significantly reduce its size, thereby decreasing the total archive footprint.

## 2.4 Decompression method

The main interest in the structure described in this article is the ability to perform targeted decompression on a query representing a subset of the compressed data. As the inverted de Bruijn graph is specifically designed for this type of operation, full decompression can be viewed as a special case of targeted decompression with the query being the entirety of the compressed data.

### 2.4.1 Targeted decompression

When a user chooses to decompress only specific input files, `kloe` first reads the entire linking file. For each listed color set matching a queried file, `kloe` retrieves the bucket’s starting position and its monochromatig sizes from the linking file. i.e. We check if the input file is present in the color set and get its associated metadata to decompress it. Using this data, `kloe` then jumps to these exact positions in the monochromatigs file and decompresses only the necessary monochromatigs into the relevant output files. Because `kloe` seeks only the required color buckets in the monochromatigs file, this method processes only relevant monochromatigs (and therefore *k*-mers) and avoids reading unnecessary data from other samples. The entire decompression workflow is described by Figure 4.

**Figure 4.**
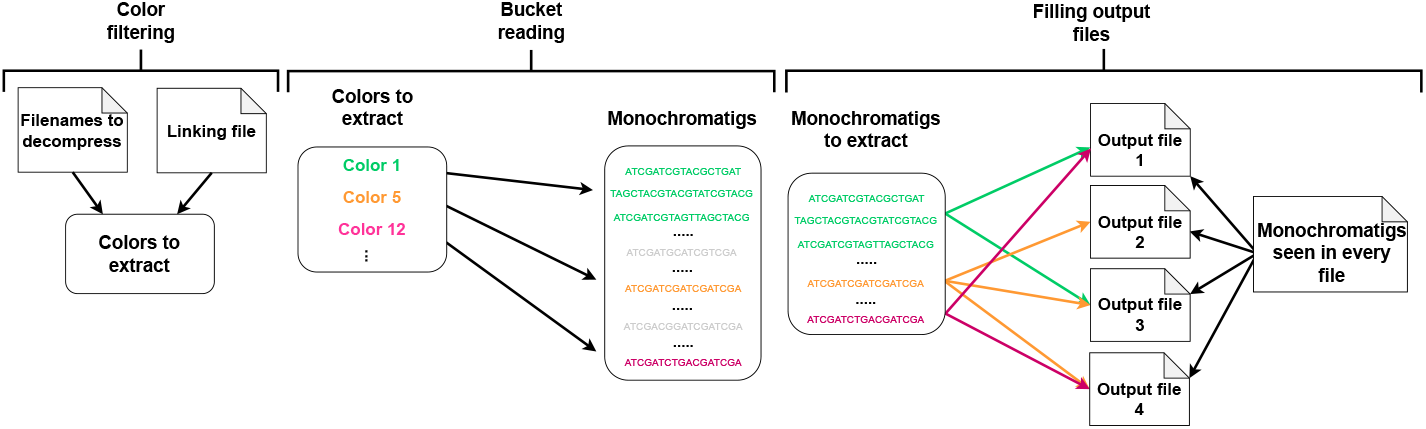
Depiction of the decompression workflow. **Color filtering** step is the gathering of necessary colors and positions of said colors in the monochromatigs file. **Bucket reading** step is reading the necessary color buckets by jumping from bucket to bucket in the monochromatigs file. **Output data** step is the writing of necessary data in every queried file (which can be the entirety of the archive or just a subset). Monochromatigs are written according to their color and the ones seen in every sets are written everywhere.

Currently, `kloe` outputs monochromatigs rather than unitigs after decompression. These can be subsequently processed by standard tools that generate unitigs from a FASTA file to revert the inputs to their state before compression with `kloe` such as `bcalm` [7], `ggcat` [9] and `cuttlefish` [15].

## 3 Results

Every experiment was run on a Supermicro Superserver SYS-2049U-TR4 with 3 TB RAM and 4 Intel SKL 6132 14-cores @ 2.6 GHz.

### 3.1 Evaluated datasets and tools

#### 3.1.1 Data

Experiments were run on 256 Human unitigs files downloaded from Logan [8]. These unitigs occupy 664 GB of space using Logan’s `zstd`-based compression system, for a mean size of 2.77 GB per `zstd`-compressed file. Lists of accessions for these data are available on `kloe`‘s Github page.

#### 3.1.2 Tools

We compared `kloe` with general-purpose compressors `xz` (efficient but slow), `zstd` (balanced speed/ratio) and `ggcat` for its compact graph-based output. Other DNA-specific tools that cannot handle unitig collections (e.g. `agc`) were not included. We did not manage to include `ESS-Color` and the *k*-mer -set compression tool proposed in [16] as we were unable to successfully run these tools on our test data.

### 3.2 Compression

#### 3.2.1 Methodology

All human unitigs were compressed from their native FASTA format. Tools were generally run with default options, a *k*-mer size of 31, and 20 threads for `ggcat` and `kloe`. To assess compression level effects, `xz` and `zstd` were also benchmarked with a fast setting (the -1 option). To ensure a fair comparison, the time to `zstd`-compress the output files was added to `ggcat`‘s graph construction runtime, accounting for its uncompressed output.

#### 3.2.2 Archive Size Comparison

Figure 5 compares tool archive sizes. Dedicated DNA compressors, `kloe` and `ggcat`, significantly outperform generic ones like `zstd` and `xz`, irrespective of settings. `kloe` achieves slightly better compression than `ggcat`. This difference arises because `kloe` outputs monochromatigs that correspond to monochromatic simplitigs, whereas `ggcat` produces monochromatic unitigs. The tested version of `ggcat` did not support colored simplitigs. Moreover, `ggcat` did not further compress its output files when handling color information (i.e., the FASTA file containing sequences and the accompanying color file). To ensure a fair comparison, we applied `zstd` compression to its indexes before measuring their sizes. In `kloe`, as collection size increases, the linking file, due to a naive representation, occupies an increasingly large fraction of the total archive. For example, in a 43 GB compressed collection of 256 human genomes, the DNA sequences themselves account for only 21 GB. Improving the compression of the linking structure could therefore provide additional space savings. Figure 5 shows compression results for each evaluated tools.

**Figure 5.**
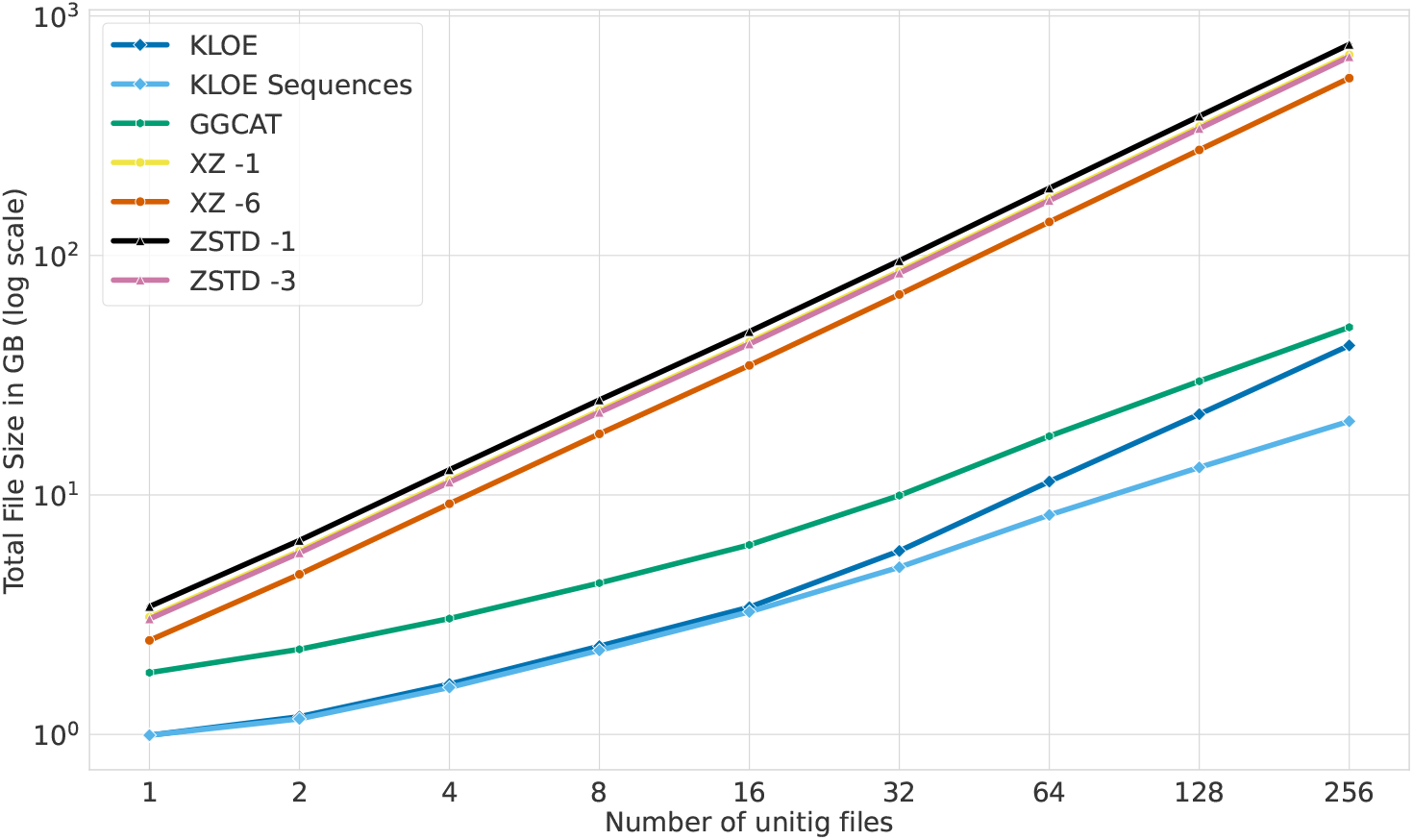
Size of compressed archives for an increasing number of unitigs from input human WGS samples. The light blue line named “`kloe` sequences” represents the disk size taken by the sole sequences in `kloe`’s compressed archive.

#### 3.2.3 RAM Usage and Compression Time

The performance profile of `kloe` is defined by its specific resource trade-offs. In terms of runtime, `kloe` is several times slower than `ggcat` and the fastest `zstd` setting. However, it remains faster than the default `zstd` and the fastest `xz` compression settings, positioning it as a viable option (Figure 6).

**Figure 6.**
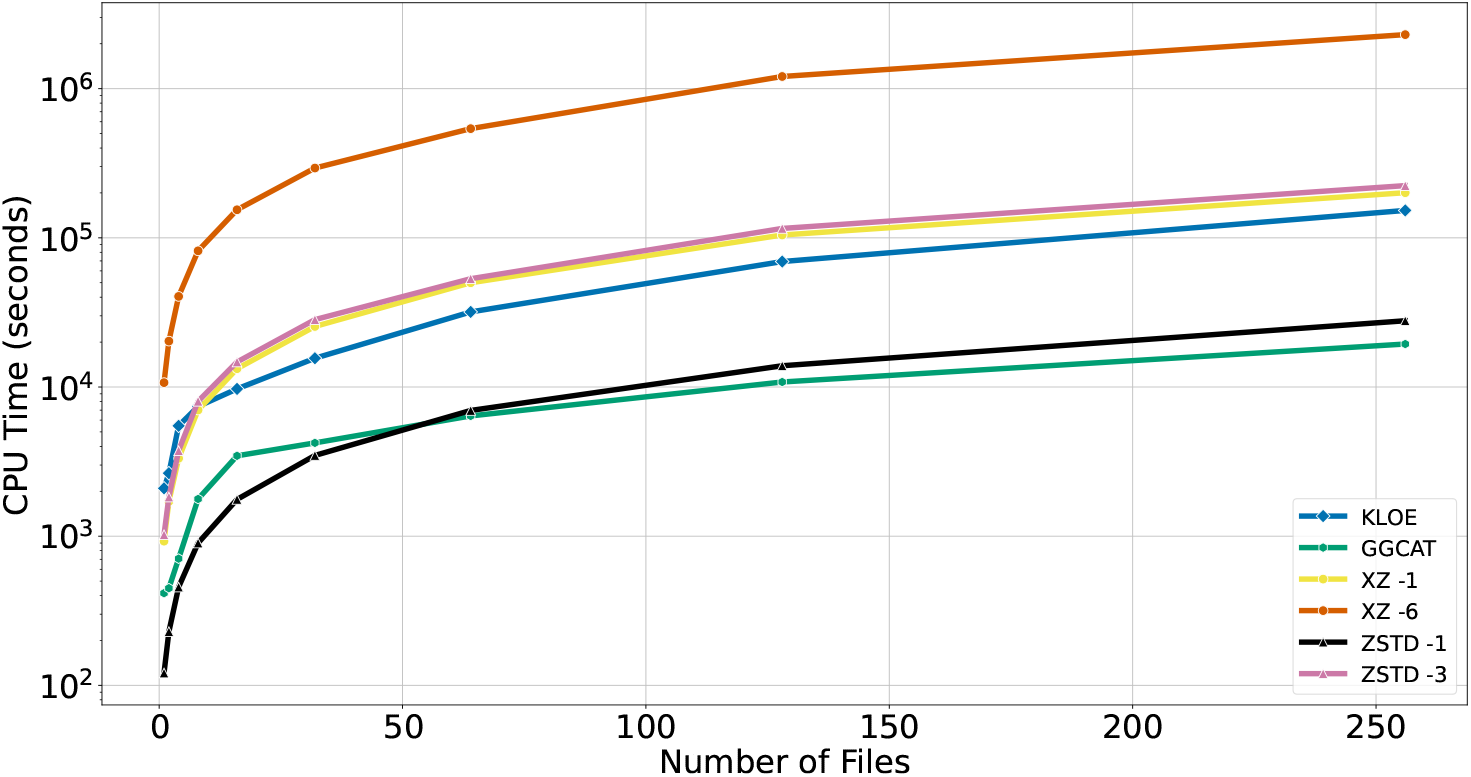
Compression time for an increasing number of unitigs from input human WGS samples.

As shown in Figure 7, the primary trade-off for `kloe` is its high memory consumption. To compress 256 human unitig files, it required over a terabyte of RAM. This approach presents a different trade-off compared to ggcat. Although ggcat uses significantly less memory at 47 gigabytes, it demands extensive disk space, with temporary files ranging from 1 to 7 terabytes. Limited-context tools `zstd` and `xz` demonstrate the lowest resource usage overall, requiring only several megabytes of RAM.

**Figure 7.**
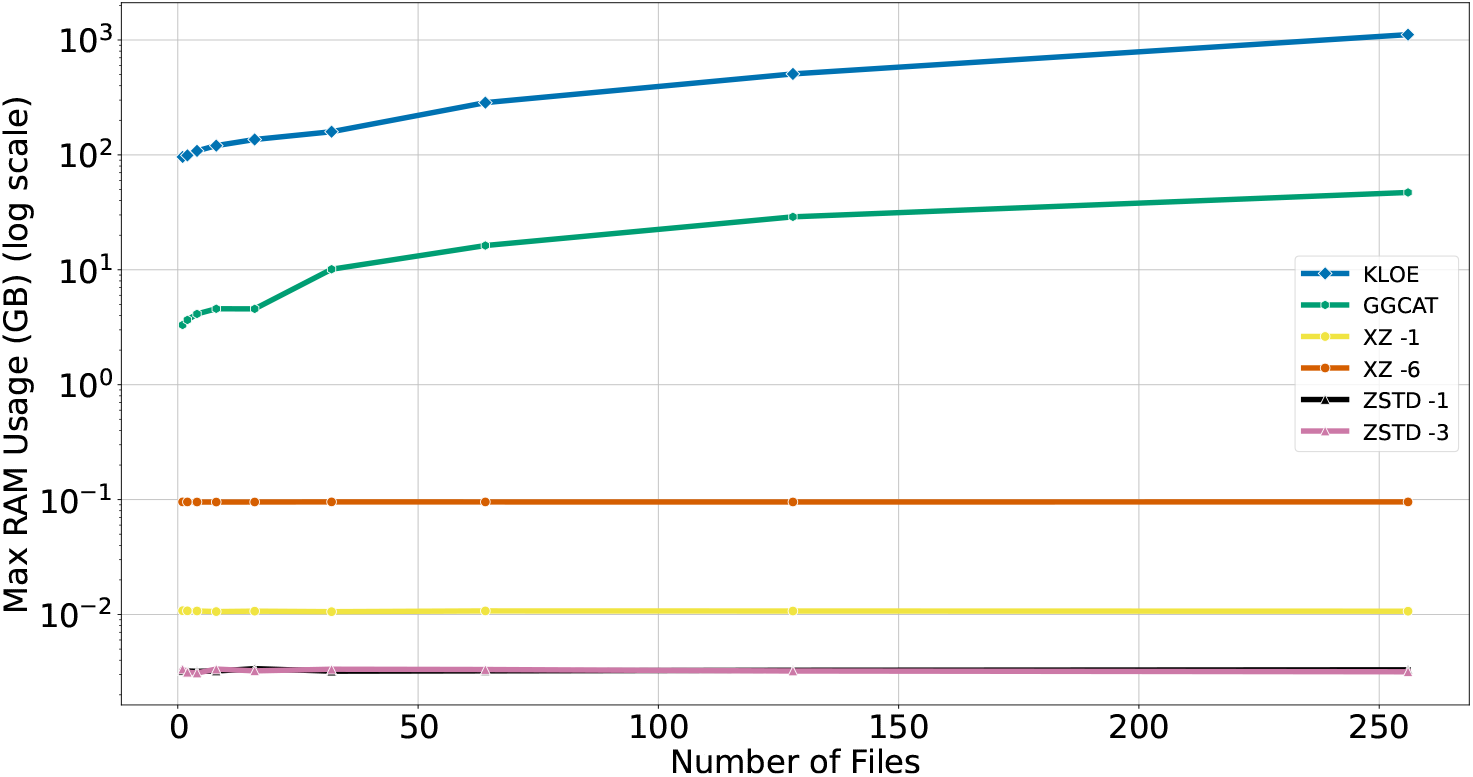
RAM usage for compression of an increasing number of unitigs from input human WGS samples.

### 3.3 Decompression

To our knowledge, `ggcat` does not provide an easy way to reconstruct the original inputs from the colored de Bruijn graph, as it was not designed for this purpose. Recovering the input sequences from the cdbg would require querying each *k*-mers for its associated colors and then assigning them back to the appropriate input files one by one. A GitHub issue addressing this limitation was posted by a user, with the author stating that such functionality is under development (https://github.com/algbio/ggcat/issues/46). However, at the time of writing, no updates have been made in this direction. Given this, we present decompression results only for `xz`, `zstd`, and `kloe`‘s performance.

#### 3.3.1 Targeted Decompression

Targeted decompression with `xz` and `zstd` is inefficient, as it requires decompressing the entire archive to filter out requested datasets. In contrast, `kloe` achieves orders-of-magnitude faster decompression by reading only the fraction of the collection needed to retrieve a given file. As a baseline, Figure 8 reports the full archive decompression times for `xz` and `zstd`. The main performance bottleneck for `kloe` is the CPU time spent parsing the linking file which prevents in practice the *O*(*S*_*j*_) complexity. Optimizing its structure could further reduce the computational cost of decompression.

**Figure 8.**
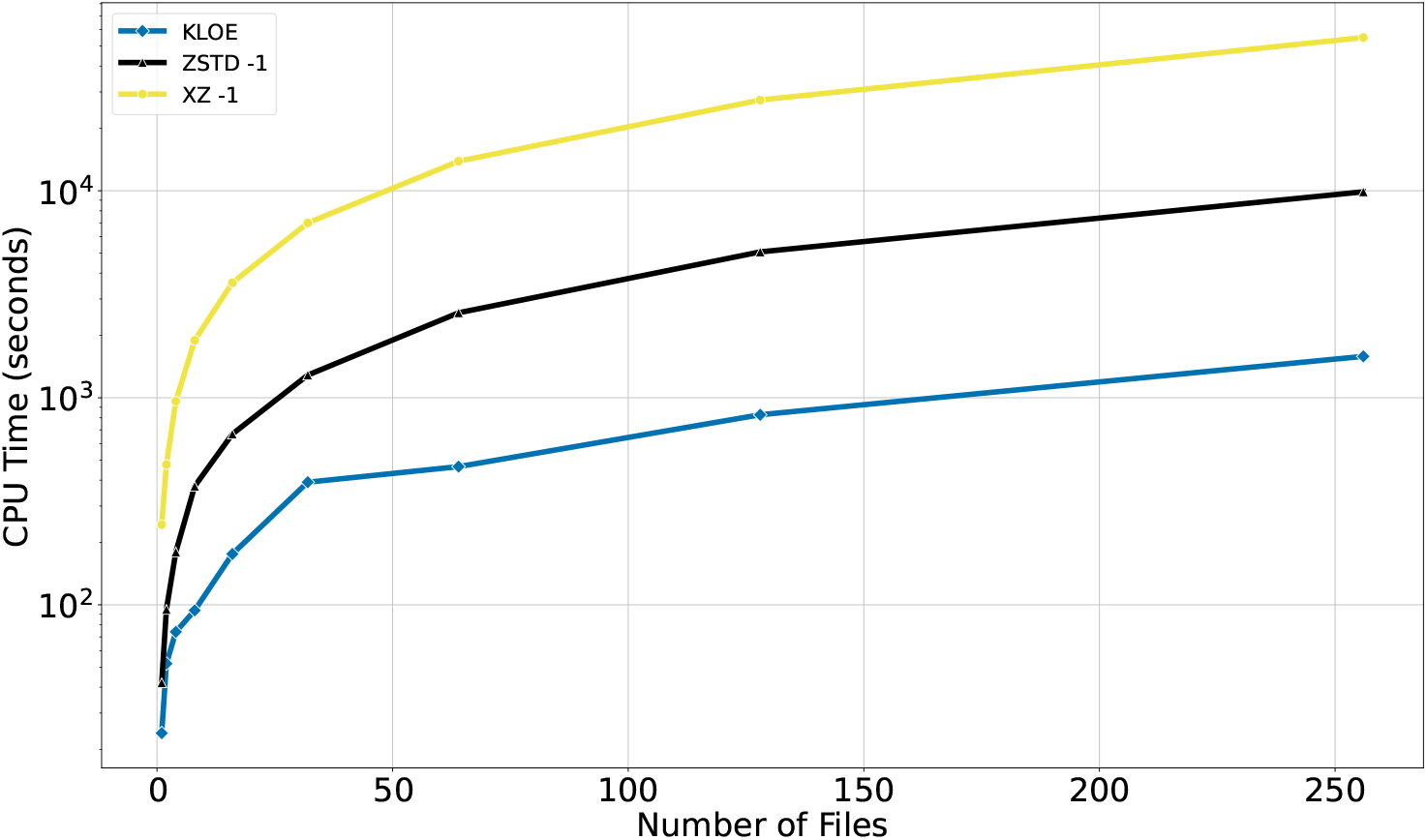
Decompression time of a single unitig file for collections containing an increasing number of unitigs from input human WGS samples.

#### 3.3.2 Full archive decompression

Performance results for the baseline operation of full dataset decompression are provided in the supplementary materials, as this is not the paper’s primary focus.

## 4 Discussion

In this work, we introduce the concept of an **inverted colored de Bruijn graph index**, a novel data structure designed to address a critical, yet overlooked, challenge in large-scale genomics: the efficient, targeted retrieval of specific datasets from a compressed collection of *k*-mer sets. The prevailing paradigm for indexing collections of genomes or sequencing experiments, the colored de Bruijn Graph (cDBG), has been extensively optimized for *k*-mer -to-color queries, that is, identifying the set of experiments containing a given *k*-mer . While invaluable for applications like differential genomics, this design is fundamentally inefficient for the inverse problem of retrieving all *k*-mers originating from a specific experiment (a *color-to-*k*-mer* query). Crucially, no existing solution is able to perform such a targeted decompression in time proportional to the size of the document to be decompressed; instead, they require complete decompression or a full scan of the archive. This presents a significant bottleneck for managing massive genomic repositories. Finding an efficient data structure capable of performing both types of queries optimally is a key open problem.

Our proof-of-concept implementation, `kloe`, is the first tool designed specifically to solve this targeted decompression challenge, demonstrating the practical feasibility and power of the inverted index concept. The core of our method lies in reorganizing the data around colors rather than *k*-mer sequences. By assembling *k*-mers into **monochromatigs**—paths of nodes that share an identical color set - and then sorting these sequences into “color buckets” on disk, `kloe` directly maps a color to the specific data blocks containing all of its associated *k*-mers . This architecture fundamentally reorients the data organization to align with the targeted retrieval query, allowing `kloe` to read only the necessary portions of the archive, bypassing all irrelevant data.

The performance of `kloe` underscores the advantages of this approach. Our experiments show that it achieves a compression ratio competitive with specialized tools like `ggcat` and far superior to general-purpose compressors. More critically, it excels at its primary design goal. Applied to 256 human unitig samples, `kloe` achieved a 15x compression ratio, reducing a 664 GB input to a 43 GB compressed file, while enabling selective retrieval of input files in under 30 minutes. This is a feat that is orders of magnitude faster than what is possible with state-of-the-art techniques, which would require full decompression.

As a proof-of-concept, `kloe` has limitations, which highlight directions for future research. The most significant trade-off in its current version is a high memory consumption during compression, a consequence of the in-memory data structures used to manage colors and build monochromatigs. Secondly, `kloe` currently does not decompress in *O*(|*s*_*j*_ |) complexity; it has to read every color identifier to target specific buckets. This might become a bottleneck with the increase of distinct colors in the archive. Future work should focus on implementing memory-efficient, disk-based algorithms for the sorting and assembly stages as well as improving complexity during decompression by optimizing metadata organisation in the archive. Furthermore, the linking file, while essential, is currently implemented with a naive structure. Optimizing its representation and compression could yield substantial reductions in the archive’s metadata overhead, further improving the overall compression ratio. Also, because of this naive organisation, the practical targeted decompression complexity is *O*(|*Z* |). Improving metadata organisation is essential in achieving practical targeted decompression complexity in *O*(|*s*_*j*_ |) as well as scalability in archive size. Finally, extending `kloe` to directly reconstruct the original unitig files (or any other SPSS), rather than outputting monochromatigs, would streamline its practical integration into existing bioinformatics pipelines.

The need for scalable solutions for storing and querying massive genomic datasets is becoming increasingly acute. We envision that `kloe` and related future works will play an essential role in compressing very large collections of unitigs, such as those from the 2.18 PB Logan dataset. By defining and providing a first solution to the problem of efficient targeted decompression, we hope that the community will be interested in this novel problem that is challenging and relevant for the future usage of massive databases.

## Acknowledgments

This work was funded by ERC CoG 101088572 (“IndexThePlanet”).

