## Supplemental Table 4 for "Inverted colored de Bruijn Graph for practical kmer sets storage"

| Nb files | KLOE | GGCAT | XZ -1 | XZ -6 | Zstd -1 | Zstd -3 |
|---|---|---|---|---|---|---|
| 1 | 95.96 GB | 3.31 GB | 11.05 MB | 97.77 MB | 3.32 MB | 3.43 MB |
| 2 | 99.05 GB | 3.66 GB | 11.02 MB | 97.71 MB | 3.30 MB | 3.22 MB |
| 4 | 108.44 GB | 4.13 GB | 10.99 MB | 97.60 MB | 3.28 MB | 3.17 MB |
| 8 | 120.40 GB | 4.58 GB | 10.86 MB | 97.52 MB | 3.30 MB | 3.43 MB |
| 16 | 136.16 GB | 4.57 GB | 10.95 MB | 97.60 MB | 3.46 MB | 3.31 MB |
| 32 | 159.09 GB | 10.11 GB | 10.84 MB | 97.80 MB | 3.29 MB | 3.42 MB |
| 64 | 284.93 GB | 16.26 GB | 11.00 MB | 97.71 MB | 3.31 MB | 3.40 MB |
| 128 | 506.73 GB | 28.90 GB | 10.98 MB | 97.66 MB | 3.35 MB | 3.30 MB |
| 256 | 1113.54 GB | 47.10 GB | 10.85 MB | 97.77 MB | 3.38 MB | 3.25 MB |

**Table 1.** Max RAM Usage for compression of a growing number of Human unitigs
