## Supplemental Table 3 for "Inverted colored de Bruijn Graph for practical kmer sets storage"

| Nb files | KLOE | GGCAT | XZ -1 | XZ -6 | Zstd -1 | Zstd -3 |
|---|---|---|---|---|---|---|
| 1 | 00:34:52 | 00:06:55 | 00:15:26 | 02:58:20 | 00:02:01 | 00:17:03 |
| 2 | 00:44:03 | 00:07:28 | 00:28:29 | 05:38:26 | 00:03:49 | 00:30:30 |
| 4 | 01:31:33 | 00:11:47 | 00:55:24 | 11:14:39 | 00:07:36 | 01:02:24 |
| 8 | 02:03:28 | 00:29:36 | 01:56:49 | 22:44:02 | 00:15:00 | 02:13:43 |
| 16 | 02:41:53 | 00:57:38 | 03:41:04 | 42:49:59 | 00:29:24 | 04:05:14 |
| 32 | 04:19:48 | 01:10:21 | 07:04:23 | 81:34:23 | 00:57:57 | 07:51:33 |
| 64 | 08:51:04 | 01:46:39 | 13:53:07 | 149:50:30 | 01:56:03 | 02:46:33 |
| 128 | 19:15:42 | 03:00:04 | 29:00:50 | 335:08:52 | 03:51:21 | 32:06:43 |
| 256 | 42:24:23 | 05:23:34 | 55:44:11 | 639:29:59 | 07:43:25 | 62:11:43 |

**Table 1.** Compression time for a growing number of Human unitigs. (HH:mm:ss)
