## Supplemental Table 2 for "Inverted colored de Bruijn Graph for practical kmer sets storage"

| Nb files | KLOE | Zstd -1 | XZ -1 |
|---|---|---|---|
| 1 | 15892 | 5668 | 3376 |
| 2 | 757804 | 5692 | 3356 |
| 4 | 1751992 | 5796 | 3352 |
| 8 | 3451508 | 5732 | 3348 |
| 16 | 6017300 | 5752 | 3360 |
| 32 | 29389904 | 5776 | 3400 |
| 64 | 93216912 | 5748 | 3376 |
| 128 | 244684248 | 5688 | 3312 |
| 256 | 661784096 | 5672 | 3360 |

**Table 1.** RAM usage for decompression of a growing number of Human unitigs compressed collections
