## Supplemental Table 1 for "Inverted colored de Bruijn Graph for practical kmer sets storage"

| Nb files | KLOE | Zstd -1 | XZ -1 |
|---|---|---|---|
| 1 | 00:00:25 | 00:00:42 | 00:04:04 |
| 2 | 00:00:52 | 00:01:35 | 00:07:56 |
| 4 | 00:01:14 | 00:03:00 | 00:16:03 |
| 8 | 00:01:34 | 00:06:14 | 00:31:35 |
| 16 | 00:02:56 | 00:11:08 | 00:59:57 |
| 32 | 00:06:31 | 00:21:27 | 01:56:33 |
| 64 | 00:07:45 | 00:42:53 | 03:51:24 |
| 128 | 00:13:48 | 01:24:25 | 07:37:08 |
| 256 | 00:26:25 | 02:44:43 | 15:15:55 |

**Table 1.** Decompression time for a queried sample among a growing number of Human unitigs compressed collections
